# Cyclopamine sensitizes Glioblastoma cells to Temozolomide treatment through Sonic Hedgehog pathway

**DOI:** 10.1101/2020.04.13.034645

**Authors:** Gabriela Basile Carballo, Diana Matias, Jessica Honorato, Luciana Santos Pessoa, Ananias Matos Arrais Neto, Tania Cristina Leite de Sampaio e Spohr

## Abstract

**Aim:** Glioblastoma is an extremely aggressive glioma, resistant to radio and chemotherapy usually performed with temozolomide. One of the main reasons for glioblastoma resistance to conventional therapies is due to the presence of cancer stem-like cells. These cells could recapitulate some signaling pathways important for embryonic development, such as Sonic hedgehog. Here, we investigated if the inhibitor of the Sonic hedgehog pathway, cyclopamine, could potentiate the temozolomide effect in cancer stem-like cells and glioblastoma cell lines *in vitro*.

**Main methods:** The viability of glioblastoma cells exposed to cyclopamine and temozolomide treatment was evaluated by using 3-(4,5-dimethylthiazol-2-yl)-2,5-diphenyltetrazolium bromide assay while the induction of apoptosis was assessed by western blot. The stemness properties of glioma cells were verified by clonogenic and differentiation assay and the expression of stem cell markers were measured by fluorescence microscopy and western blot.

**Key findings:** The glioblastoma viability was reduced by cyclopamine treatment. Cyclopamine potentiated temozolomide treatment in glioblastoma cell lines by inducing apoptosis through activation of caspase-3 cleaved. Conversely, the combined treatment of cyclopamine and temozolomide potentiated the stemness properties of glioblastoma cells by inducing the expression of SOX-2 and OCT-4.

**Significance:** Cyclopamine plays an effect on glioblastoma cell lines but also sensibilize them to temozolomide treatment. Thus, first-line treatment with Sonic hedgehog inhibitor followed by temozolomide could be used as a new therapeutic strategy for glioblastoma patients.

## 1. Introduction

Glioblastoma (GBM) is the most common primary brain tumor [1]. The median survival of patients diagnosed with GBM is nearly 15 months even after conventional treatment by surgery, radiotherapy, and chemotherapy with temozolomide (TMZ) [2–4].

GBM radio and chemoresistance is related to several mechanisms, such as the presence of glioma stem cells (GSCs), for instance [5]. Some studies have shown that TMZ is inefficient in inducing GSC death [6,7]. Moreover, the tumor microenvironment (TME) recapitulates the activation of several pathways including Sonic hedgehog (SHH), *Wingless* (WNT), Notch and transforming growth beta (TGF-β) [8–11], responsible for GSCs’ maintenance in the tumor mass [12,13]. In fact, the upregulation of the SHH pathway preserves the main characteristics of GSC such as the ability of self-renewal and chemoresistance in tumors [14,15]. In general, the SHH signaling pathway is activated when SHH binds to 12-transmembrane protein Patched (PTCH), named PTCH1 or PTCH2 and consequently occurs the releasing of 7-transmembrane protein Smoothened (SMO), which in turn can be activated and stabilized initiating the SHH downstream signaling cascade [11,16–18]. This downstream cascade produces the translocation of GLI family proteins from the cytoplasm to the nucleus and consequently, it starts the transcription of target genes including *PTCH, GLI, SOX-2*, and *OCT-4* [19,20]. Cyclopamine, isolated from *Veratrum californicum* plant, is a natural inhibitor of SHH pathway by binding to SMO and consequently blocks the downstream signaling pathway [11,21–24].

It has already been demonstrated that high levels of Gli1 are associated with upregulation of O-6-methylguanine-DNA methyltransferase (MGMT) and consequently induces the glioma chemoresistance to TMZ [25]. Therefore the inhibition of the SHH pathway could be a promising therapeutic approach by restoring the chemosensitivity of GBM cells to TMZ. Here, we analyzed if the inhibition of the SHH pathway with cyclopamine could potentiate the TMZ effect in different GBM cell lines *in vitro*.

## 2. Material and methods

### 2.1. Reagents

All culture reagents as well as the secondary antibodies, conjugated to Alexa Fluor 488, 546 and HRP, Fluoromount-G, BCA™ Protein Assay kit, Super Signal™ West Pico or West Femto Chemiluminescent Substrate, Trizol, oligodT (12–18) primer, High-Capacity cDNA Reverse Transcription Kit and Power Sybr Green Master Mix, were obtained from Life Technologies (Carlsbad, CA, USA). The 4-6-diamino-2-phenylindole (DAPI), 3-(4, 5-dimethylthiazol-2-yl)-2,5-diphenyl tetrazolium bromide (MTT), TMZ, anti-α-tubulin and dimethyl sulfoxide (DMSO) were purchased from Sigma-Aldrich Corp. (St. Louis, MO, USA). Anti-SOX-2, anti-OCT-4 and anti-caspase 3 antibodies were obtained from Cell Signaling Technology (Danvers, MA, USA). Hybond-P polyvinylidene difluoride (PVDF) transfer membrane was obtained from BioRad Laboratories (Berkeley, CA, USA). Anti-PTCH1, anti-SHH, anti-GAPDH were obtained from Abcam (Cambridge, MA, USA). Cyclopamine was obtained from Toronto Research Chemicals (Toronto, CA). Polyclonal Rabbit Anti-GFAP was obtained from Dako (Glostrup, Denmark).

### 2.2. Maintenance of Cell Line Cultures

All the human GBM cell lines used (GBM95, GBM02 and, GBM03) were established and characterized in our laboratory as previously described [26,27]. Foreskin fibroblast cell line nh-skp-FB0012 was obtained from Rio de Janeiro Cell bank. These cells were maintained in Dulbecco’s Modified Eagle Medium/Nutrient Mixture F-12 (DMEM/F12), supplemented with 10% of fetal bovine serum (FBS), and maintained at 37°C in an atmosphere containing 95% air and 5% CO_2_. For the assays performed using glioma stem cells (GSCs), the cells were maintained in culture according to our previous studies [5]. The cells were plated in a defined NS34 serum-free medium supplemented with sodium pyruvate, glutamine, B-27, G-5 and, N-2 supplements, penicillin and streptomycin [5] to maintain the GSCs properties.

### 2.3. MTT assay

The GBM 95, GBM02, GBM 03 and foreskin fibroblast cells were plated into 96 well plate at 5% FBS. The cell viability was assessed by the MTT reduction colorimetric assay. Cell viability was measured at different conditions of treatment: i) 192 hours of incubation with three different concentrations of cyclopamine (5 μM, 7.5 μM and 10 μM); ii) 192 hours after cyclopamine treatment, the cells were maintained in culture for more 192 hours in the absence or presence of cyclopamine - we defined as ‘long term MTT’ (in a total time of 16 days, 384 hours); iii) 144 hours (6 days) of incubation with temozolomide at different concentrations (100 μM, 200 μM, 400 μM, 600 μM); iv) 144 hours of cyclopamine 7.5 μM and TMZ 250 μM treatment. MTT was added to each well for 2 hours and blue formazan crystals produced were dissolved by adding 100 μL of DMSO. Absorbances were measured at 570 nm.

### 2.4. Immunofluorescence

GBM cells were cultured on coverslips in 24-well plates, as previously described [28]. Briefly, cultured cells were fixed with 4% paraformaldehyde (PFA) for 30 min and permeabilized with 0.1% Triton X-100 for 5 minutes at room temperature. The cells were blocked with 5% bovine serum albumin (BSA) for 1 hour at room temperature. GSCs were fixed as described above; however, the permeabilization step was performed by using 0.3% of Triton X-100 for 20 minutes and then blocked with 5% BSA for 1 hour. The fixed cells were incubated overnight at 4°C with rabbit anti-SOX-2 (1:400) and with rabbit anti-OCT-4 (1:400) primary antibodies. Then, after the cells were washed with PBS and secondary antibodies staining were performed using goat anti-rabbit IgG conjugated with Alexa fluor 488 and 546 (1:400) for 2 h, at room temperature. Thereafter, the cells were washed with PBS, stained with DAPI and mounted in Fluoromount-G. Cells were imaged using a DMi8 advanced fluorescence microscope (Leica Microsystems, Germany) and analyzed with Leica LAS X Lite. Images were processed using the software ImageJ 1.49v (Wayne Rasband, National Institutes of Health).

### 2.5. Western blotting

GBM cells were plated into 6-well plates and incubated for each condition as previously described. The lysis buffer was added (RIPA 1x: NaCl, EDTA, Triton X-100, Sodium deoxycholate, SDS, di-water, 1M Tris-HCl, pH 7,6) for 30 minutes and the cell lysates were sonicated and centrifuged at 4°C, 20817 g for 10 minutes. The supernatants were analyzed for protein content by the BCA™ Protein Assay kit. 30 µg or 70 µg of protein per lane was electrophoretically separated in 8%, 10% and, 12% sodium dodecyl sulfate-polyacrylamide gel. Western blotting was performed as described by Towbin et al. 1992 [29] with minor modifications introduced by [30], 2014). After separation, the proteins were electrically transferred to a PVDF transfer membrane. The PVDF membranes were then blocked with 5% non-fat milk in Tris-buffered saline with 0.1% Tween-20 (TBS-T) for 1 hour, incubated with specific primary antibodies overnight at 4 °C. Thereafter, the membranes were washed with TBS-T and incubated with peroxidase-conjugated antibodies. The signals of SOX-2 (1:1000), OCT-4 (1:1000), PTCH1 (1:1000), SHH (1:1000), CASPASE 3, GAPDH

(1:2000), and α-TUBULIN (1:4000) were detected using SuperSignal™ West Pico or West Femto Chemiluminescent Substrate chemiluminescence in ChemiDoc MP imaging system (BioRad, Berkeley, USA). The densitometry analyses were performed using Bio-Lab Software, (Bio-Rad, Berkeley, USA), and the values obtained represent the ratio between the immunodetected protein and the loading control (GAPDH or α-Tubulin).

### 2.6. Clonogenic Assay

In order to evaluate the clonogenic capability of GBM cells, a single GBM cell was seeded with 100 μL of NS34 medium in each well of a 96-well plate. The cells were maintained in the NS34 medium for 4 weeks by replacing the stem cell medium every 2 days. The colonies/oncospheres formations were counted every week by observation under a phase-contrast microscope (DMi1, Leica Microsystems, Germany).

### 2.7. Differentiation assay

The differentiation assay was performed after 6 days of treatment with cyclopamine 7.5 μM and TMZ 250 μM under different established conditions. GSCs were isolated from GBM treated cells in the presence of NS34 medium in a 24-wells plate. After 2 weeks, the formation of floating oncospheres was observed, typical characteristics of GSCs. Thereafter, the NS34 medium was replaced by DMEM/F-12 supplemented with 2% FBS to promote the differentiation of GSCs for 10 days. Cells were fixed with 4% PFA and immunofluorescence for anti-GFAP (1:500) was performed as described above, since GSCs-differentiated express GFAP. Moreover, the morphological features of GSCs in the presence of FBS, a monolayer cell growth, were observed.

### 2.8. qRT-PCR

Total RNA was extracted from approximately 10^6^ cells via the Maxwell 16 LEV simply RNA Tissue Kit (Cat. #AS1280, Promega, Madison, WI) on a Maxwell 16 Instrument (Cat. #AS2000, Promega, Madison, WI) according to the manufacturer’s instructions. After the extraction the samples were submitted to a reverse transcription reaction using High Capacity cDNA Reverse Transcription Kit (Applied Biosystems, California, USA) according to manufacturer’s instructions. cDNA concentration was measured with a NanoDrop Lite Spectrophotometer (Thermo Fisher Scientific, Massachusetts, USA). Quantitative polymerase chain reactions (qPCR) were carried out in triplicate using 50 ng cDNA, SsoAdvanced Universal SYBR Green Supermix (BioRad, California, USA) following the manufacturer’s instructions, MGMT primers for the exon 4 region (Forward: 5’GTGTACCGCTCCAAGGACAA3’; Reverse: 5’GCCTTCCCTCTCCCTCTGTA3’) and GAPDH as the endogenous control (Forward: 5’GAGTCAACGGATTTGGTCGT3’; Reverse: 5’ TTGATTTTGGAGGGATCTCG 3’). To calculate relative fold variations in mRNA expression, the 2-ΔΔCT method was used and the data were analyzed using Student’s t-test. Thermal cycling was always carried out using the conditions recommended by the manufacturer in a CFX96 Touch Real-Time PCR Detector (BioRad BioRad, California, USA).

### 2.9. Coefficient of drug interaction (CDI)

The nature of drug interaction was analyzed through the coefficient of drug interaction (CDI) [31], considering CDI = AB / (A× B). According to the absorbance of each group calculated by MTT assay, AB is the ratio of the drugs combination group to the control; and A or B is the ratio of the single-agent group to the control. Thus, a CDI value of <1, = 1, or >1 indicates if the drugs are synergistic, additive, or antagonistic, respectively.

### 2.10. Quantitative analysis

All values were expressed as mean ± SD. The groups were compared by means of a one-way ANOVA test or two-way ANOVA test, Dunnet’s test, and Bonferroni probabilities, with a significance threshold of p < 0.05. All statistical analysis was performed using GraphPad Prism 6 (GraphPad Software Inc., San Diego, CA, USA). The experiments were done in triplicate, and each result represents the mean of three independent experiments.

## 3. Results

### 3.1. SHH pathway inhibition with cyclopamine interferes with GBM cell viability

As it is already known, the SHH signaling pathway is usually expressed during tumorigenesis [8–10,32]. In order to analyze if the SHH pathway is involved in cellular viability in our GBM cell lines, we performed the MTT assay in GBM95, GBM02, and GBM03 cells treated for 8 days at three different concentrations of cyclopamine (5 µM; 7.5 µM, and 10 µM). GBM cell lines presented a decrease in cell viability, especially at 7.5 µM and 10 µM concentrations (**Fig. 1 A, B, and C, and Supplementary Table 1**). No effect was observed in healthy human foreskin fibroblast cells treated with cyclopamine at the same concentrations (**Fig 1 D**). This result suggests that cyclopamine has a specific effect on the viability of tumor cell lines and the concentrations used were not toxic for non-tumor cells. To confirm the reduction observed in MTT assay, the viable cells were counted after cyclopamine treatment (**Supplementary Fig 1 C**).

**Figure 1:**
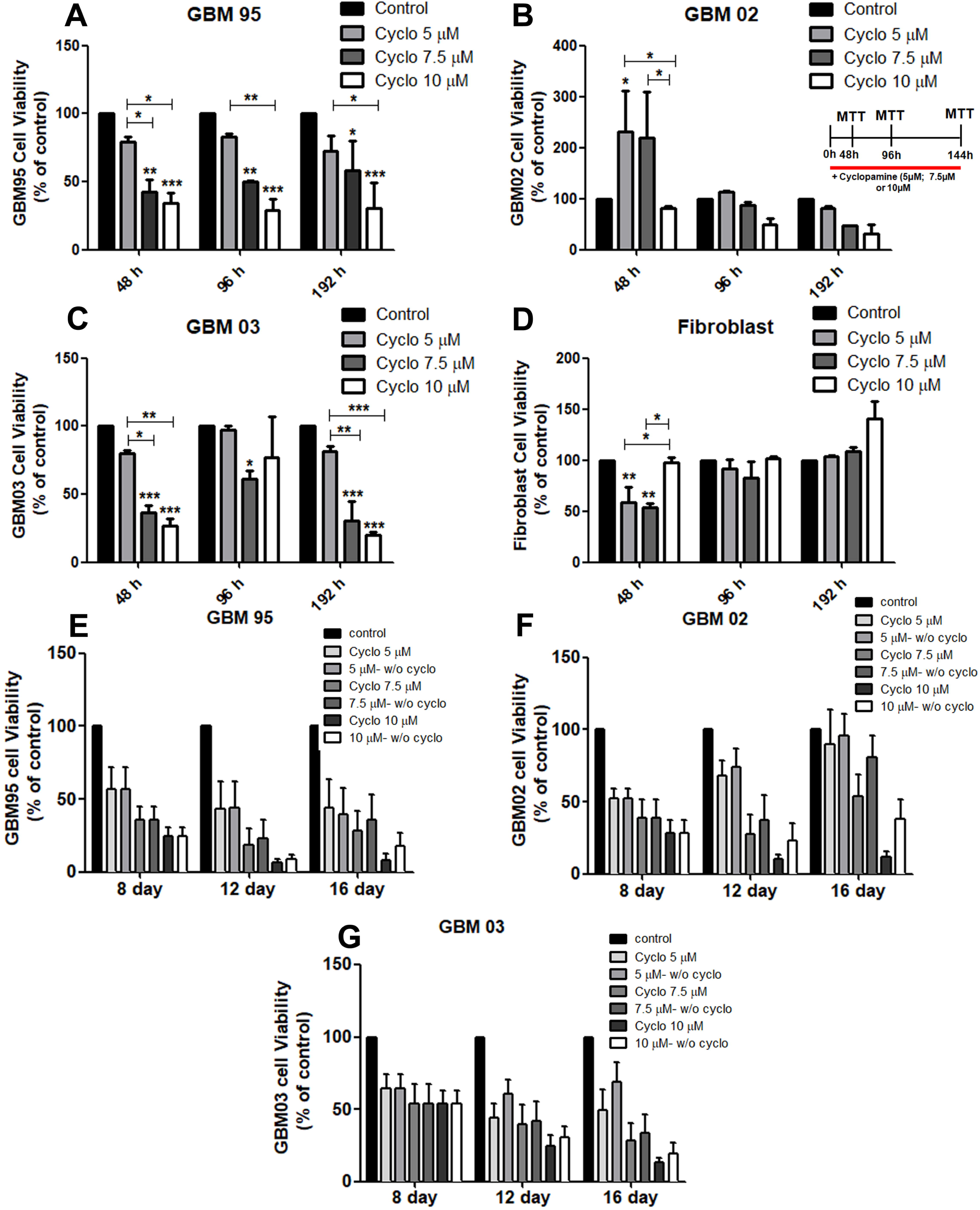
Cyclopamine interferes with GBM cell viability by MTT assay. GBM95 (**A**), GBM02 (**B**) GBM03 (**C**) and fibroblast cells (**D**) were treated with different concentrations of cyclopamine, 5μM, 7.5μM, and 10μM for 48, 96, and 192 hours (8 days). Cyclopamine induces a reduction in GBM cell viability in a dose-dependent manner; however, the non-tumor cells were not affected by cyclopamine treatment. In all GBMs cells, there was observed a presence of resistant cyclopamine-treated cells. The cellular viability was verified in GBM cells at different times after the removal of cyclopamine. None of the GBMs cell lines show an increase in viability (**H-J**). All the mean values are derived from three independent experiments. The decrease in cell viability was analyzed comparing all conditions to the control condition. (*P< 0.05; **P< 0.01 and ***P< 0.001).

We also performed an MTT assay that we referred to as ‘long term MTT’ during 384 hours (in a total time of 16 days). After 192 hours of cyclopamine treatment (8 days), the cells were maintained in culture until 16 days in the absence or presence of cyclopamine. In these conditions, none of the GBM cells presented an increase in cell viability, suggesting that even with the withdrawal of cyclopamine, the effect was sustained (**Fig 1 E, F and G and Supplementary Table 2**).

### 3.2. Cyclopamine potentiates TMZ effect in GBM cell viability

TMZ is considered the gold standard therapy for patients diagnosed with GBM [3]. In order to establish the TMZ concentration to use in combination treatments in GBM cell lines, the MTT assay was performed (**Supplemental Fig 1 and Supplementary Table 3**). The GBM cell lines were treated with four different concentrations of TMZ (100 µM, 200 µM, 400 µM, and 600 µM) for 72 and 144 hours. After 72 hours of treatment only the GBM03 showed a relative decrease in cell viability. However, after 144 hours of TMZ treatment, all GBM cell lines showed a decrease in cell viability for higher concentrations of 200 µM (**Supplemental Fig 1 A and B**).

In an attempt to evaluate if cyclopamine could potentiate TMZ effect in GBM cells, we combined both compounds in different conditions of treatment and analyzed the cell viability by using MTT assay. GBM cells were treated with monotherapy and a combination of cyclopamine 7.5µM and TMZ 250µM for 48, 72, and 144 hours (6 days). We observed that GBM cells treated with combinations of cyclopamine and TMZ (TMZ 250µM+Cyclo 7.5µM), the cell viability was reduced after 144 hours of treatment (**Fig 2 A, B, and C, and Supplementary Table 3**). We also established two conditions of treatment that we called ‘exchange condition’. In the ‘exchange condition’, the GBM cells were first submitted to cyclopamine and then to TMZ or vice-versa for 48 or 72 hours. We observed that the GBM cells submitted first to cyclopamine and then to TMZ, the cell viability was reduced (**Fig. 2 D, E, and F**). Instead, in the GBM cells first treated with TMZ followed to cyclopamine, we observed a more prominent decrease in cell viability (**Fig. 2 D, E, and F**). To corroborate the cell viability reduction, the number of cells stained with DAPI were counted in all treatment conditions. We observed a reduction in the total number of cells left in GBM cells submitted to the combination treatment (**Supplemental Fig 1 D and Supplementary Table 4**). Moreover, we calculated the coefficient of drug interactions (CDI) as according to Li and collaborators (2016) as represented in **table 1**, there was a synergistic effect in the cells treated concomitantly with TMZ 250µM+Cyclo 7.5µM during 144 hours (CDI<1) [31].

**Figure 2:**
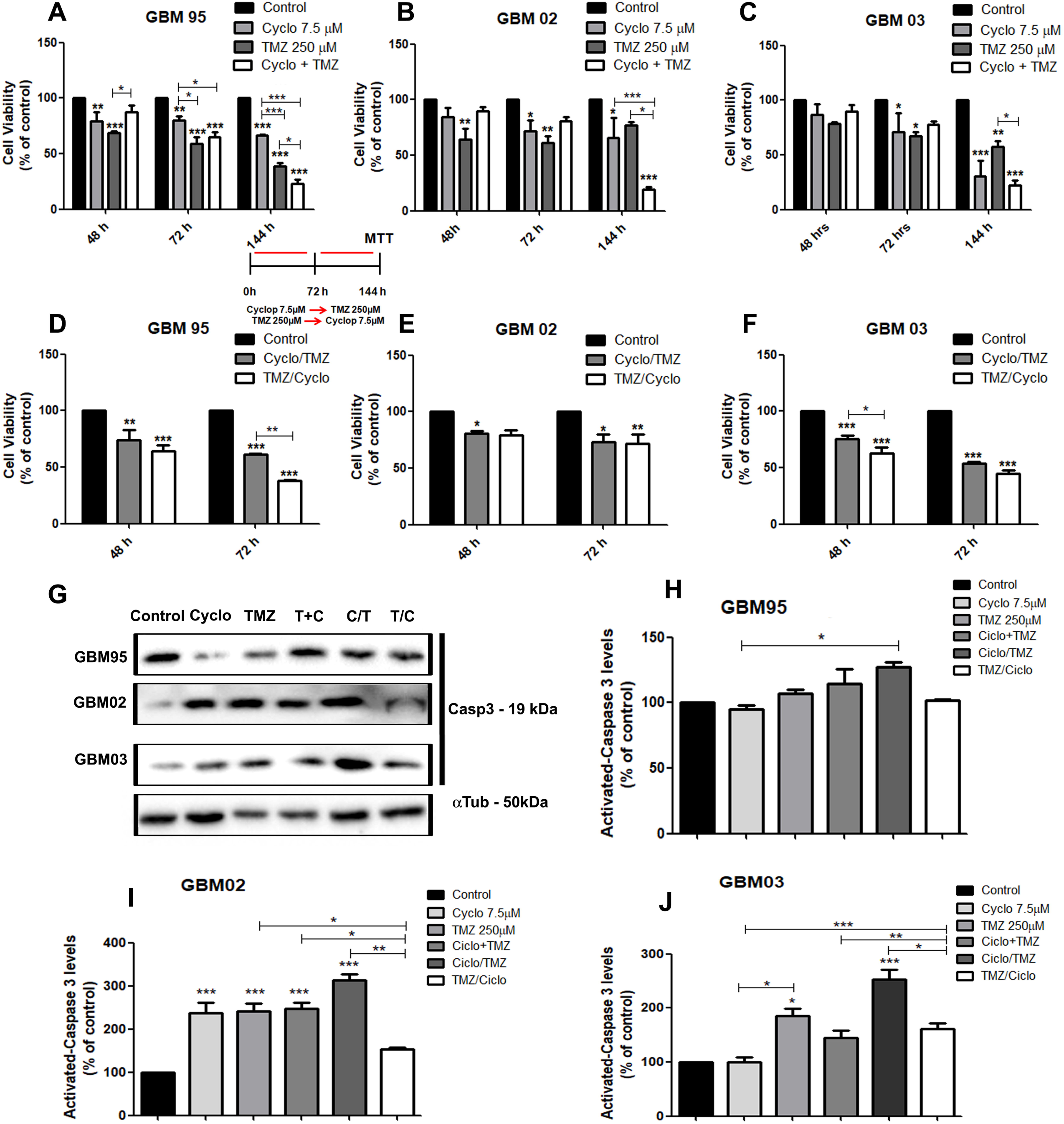
Cyclopamine potentiates TMZ effect in GBM cell viability. An MTT assay was performed to assess the cell viability of GBM95 (**A**), GBM02 (**B**), and GBM03 (**C**) cells treated with monotherapy with cyclopamine (7.5 µM) or TMZ (TMZ 250 µM) and the combination of cyclopamine (7.5 µM) with TMZ (250 µM) for 48, 72 and 144 hours (6 days). The GBM95 (**D**), GBM02 (**E**), and GBM03 (**F**) cells were submitted to ‘exchange condition’ treatment with cyclopamine (7.5 µM) or TMZ (TMZ 250 µM) for 48 and 72 hours. In the ‘exchange condition’ treatment, the cells treated with TMZ following cyclopamine treatment (TMZ/Cyclo) had a significant reduction in cell viability as compared to the control condition. Western blotting was used to analyze the protein expression of activated caspase 3 in all GBM cell lines treated with monotherapy (cyclopamine or TMZ) and different combination treatments of cyclopamine with TMZ (**G**). The α-tubulin was used as internal control. The analysis of activated caspase 3 protein levels in GBM95, GBM02, and GBM03 are represented in **H, I**, and **J**, respectively. All gels were submitted to the same assay conditions and cropped blots represent at least three independent experiments. ‘Exchange condition’ where cells were first treated with cyclopamine showed a higher expression of activated caspase 3. All the mean values were derived from three independent experiments. (*P< 0.05; **P< 0.01 and ***P< 0.001).

Taking into account the reduction of the cell viability after cyclopamine and TMZ treatment, we also analyzed the expression of activated caspase 3 that has been associated with apoptosis [33]. Our results showed that the activated caspase 3 levels were higher in GBM cells treated with cyclopamine and then with TMZ as compared to the cells submitted first to TMZ (**Fig 2 G-H and Supplementary Table 4**).

### 3.3. The combined treatment with cyclopamine and TMZ increases the stemness of GBM cells

Since the presence of GSCs is known to be one of the main reasons for the resistance to radio- and chemotherapy [34], we hypothesized that the remaining cells after the combined treatment could have acquired stem cell properties. GSCs share similar characteristics as stem cells by expressing pluripotent genes, such as SOX-2 or OCT-4, transcription factors extremely important for the maintenance of stemness of the cells [5,35], ability to form clones and floating spheres, for instance [5]. To test our hypothesis, the nuclear expression of SOX-2 and OCT-4 in those cells was evaluated. We observed that the combination treatment induces higher expression of these factors. GBM95 cells have increased SOX-2 expression in the ‘exchange condition’ cells treated first with TMZ (TMZ/Cyclo) (**Fig 3 A and G, and Supplementary Table 4**). On the other hand, in GBM95 cells first submitted to cyclopamine, we detected an increased expression of OCT-4 (Cyclo/TMZ) (**Fig 3 D and G**). In the case of the GBM02 cell line, we observed increased levels of SOX-2 and OCT-4 in both ‘exchange conditions’ (TMZ/Cyclo and Cyclo/TMZ) (**Fig 3 B, H and Supplementary Table 4**). However, the GBM03 cell line presented high levels of SOX-2 after the combination of TMZ and cyclopamine (TMZ+Cyclo). Instead of this, in the cells submitted to the ‘exchange condition’ treated first with TMZ, the OCT-4 expression was higher (TMZ/Cyclo) (**Fig 3 C, I and Supplementary Table 4**). Our results were corroborated by the analysis of the expression of these proteins by western blotting (**Fig. 4 A-F, and Supplementary Table 4)**. Interestingly, we also observed increased levels of PTCH1 and SHH in the GBM cells treated. In GBM95 cells, there was an increase of PTCH1 and SHH expression in the ‘exchange condition’ where cyclopamine was first applied (Cyclo/TMZ) (**Fig. 4 G, H and Supplementary Table 4**). In GBM02 and GBM03 cell lines in ‘exchange condition’ where TMZ was added before cyclopamine, we observed an increase of PTCH1 and SHH expression (Cyclo/TMZ) (**Fig. 4 I-L and Supplementary Table 4**).

**Figure 3:**
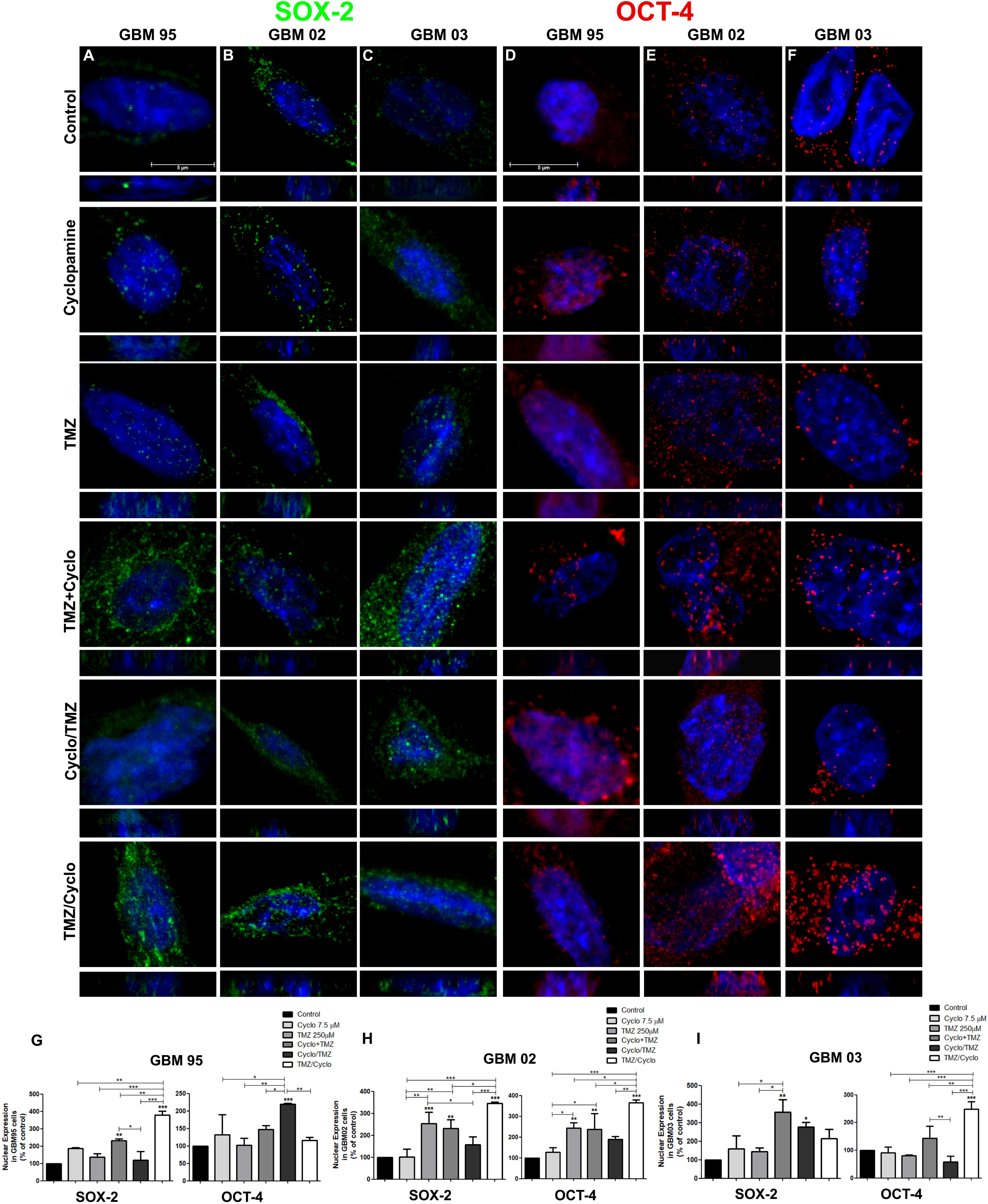
GBM cells treated with a combination of Cyclopamine and TMZ present nuclear expression of SOX-2 and OCT-4. The nuclear expressions of SOX-2 (green) and OCT-4 (red) were observed in GBM cells after monotherapy of cyclopamine and TMZ or at different combinations of cyclopamine with TMZ by immunofluorescence. Group **A** and group **D** represent the nuclear expression of SOX-2 and OCT-4 for the GBM95 cell line after 144 hours (6 days) of treatment; group **B** and **E** represent GBM02, and group **C** and **F** represent GBM03. The cell nuclei were stained with DAPI (blue). The analysis of immunofluorescence of SOX-2 and for OCT4 for GBM cell lines are represented in **G, H, I** as compared to the control conditions. The TMZ/Cyclo condition in GBM05 and GBM02 cells have increased levels of SOX-2; however, SOX-2 levels increase in GBM03 when treated with Cyclo+TMZ. The OCT-4 levels increased in GBM02 and GBM03 cells when treated with TMZ/Cyclo, whereas, in the GBM95 cells, increased levels of OCT-4 were observed in Cyclo/TMZ condition. All the mean values were derived from four independent experiments (*P< 0.05; **P< 0.01 and ***P< 0.001). Scale bar corresponds to 5 µm.

**Figure 4:**
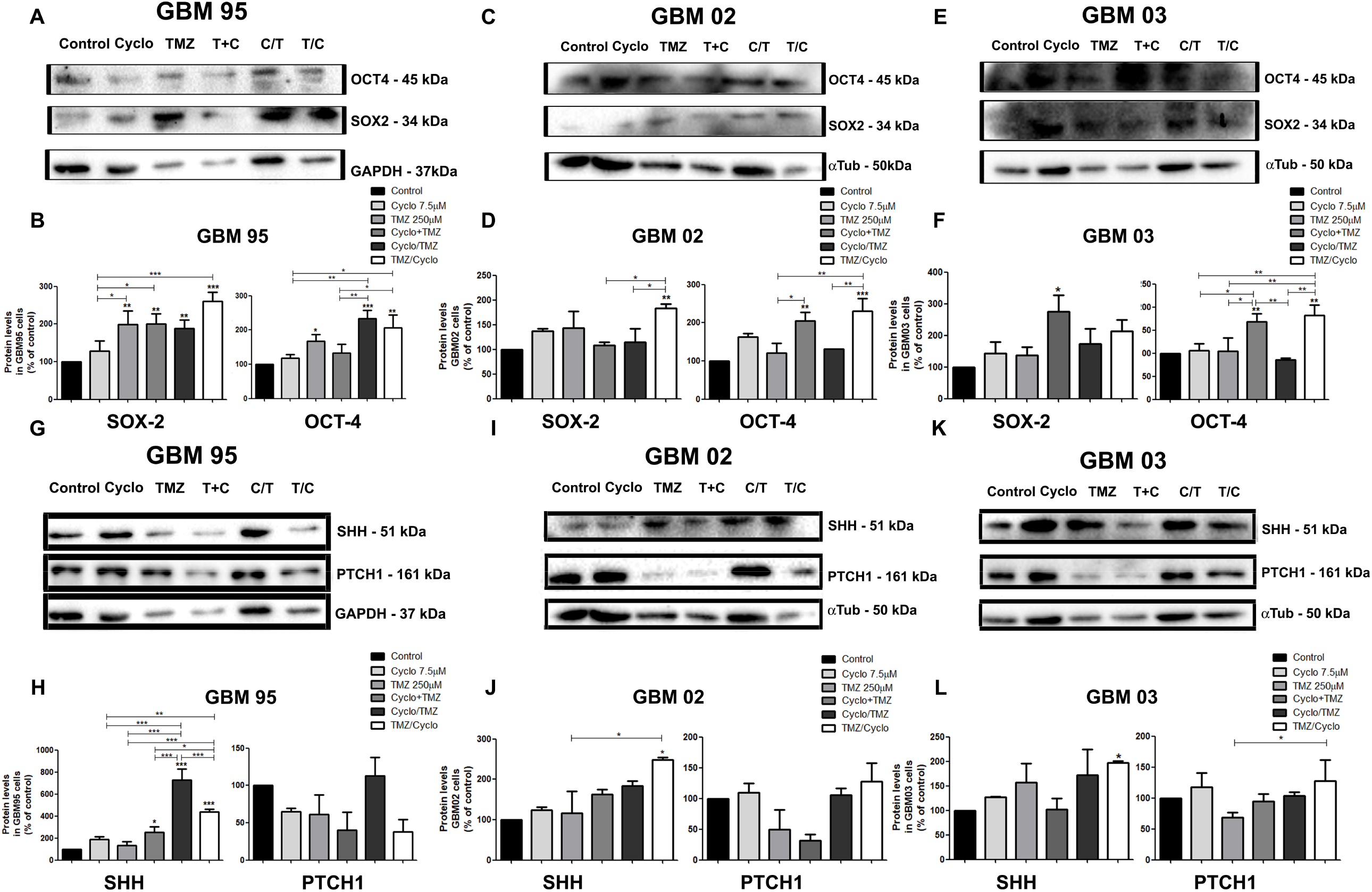
The cyclopamine and TMZ combination modulates the expression of SOX-2, OCT-4, PTCH1 and, SHH in GBM cells. The protein expression levels of SOX-2, OCT-4, PTCH1, and SHH in GBM95, GBM02, and GBM03 treated with monotherapy of cyclopamine and TMZ or in different combinations of cyclopamine with TMZ were evaluated by western blotting. **A** to **F** represent the densitometry analysis of SOX-2 and OCT-4 levels in GBM treated cells as compared to control conditions. The densitometry of PTCH1 and SHH levels in GBM treated cells are represented in **G** to **L.** GAPDH and α-tubulin were used as the internal control. All gels were submitted to the same assay conditions and cropped blots represent at least three independent experiments. All the mean values were derived from three independent experiments (*P< 0.05; **P< 0.01 and ***P< 0.001).

### 3.4. The GBM cells non-sensitive to combined treatment of cyclopamine and TMZ present CSC characteristics

To further confirm if the remaining cells after the combined treatment were GSCs, we treated the GBM cell lines with ‘exchange conditions’ for 144 hours. After that, we cells were cultured for 2 weeks in the defined NS34 medium in order to select their GSCs population. One of the main characteristics of GSCs is the ability to form floating spheres [36]. Therefore, we observed that all GBM cell lines cultured in NS34 presented floatings spheres (**Supplemental Fig. 2**). Interestingly, the GBM03 cell line presented a higher number of oncospheres as compared to GBM95 and GBM02 cells (**Fig 2S K-N**). Moreover, the cells treated were counted by DAPI staining and we observed that GBM03 cells indeed presented a higher number of cells when compared to GBM95 and GBM02 (**Fig 2S E, J, and O, and Supplementary Table 5**).

In order to corroborate the presence of GSCs, we performed a clonogenic assay. All three cell lines formed clones in all treatments (**Fig 5**). In the ‘exchange condition’, the GBM cells first treated with cyclopamine presented a higher number of clones as compared to the other two conditions (Cyclo+TMZ and TMZ/Cyclo). Moreover, when we compared only the control condition of the three cell lines, we observed, as expected, that the GBM03 cells presented a higher number of clones (**Fig 5 E, K, and Q**). We also measured the size of the oncospheres and, as expected, the GBM03 cell line also presented bigger oncospheres as compared to GBM95 and GBM02 (**Fig 5 F, L, and R, and Supplementary Table 5**).

**Figure 5:**
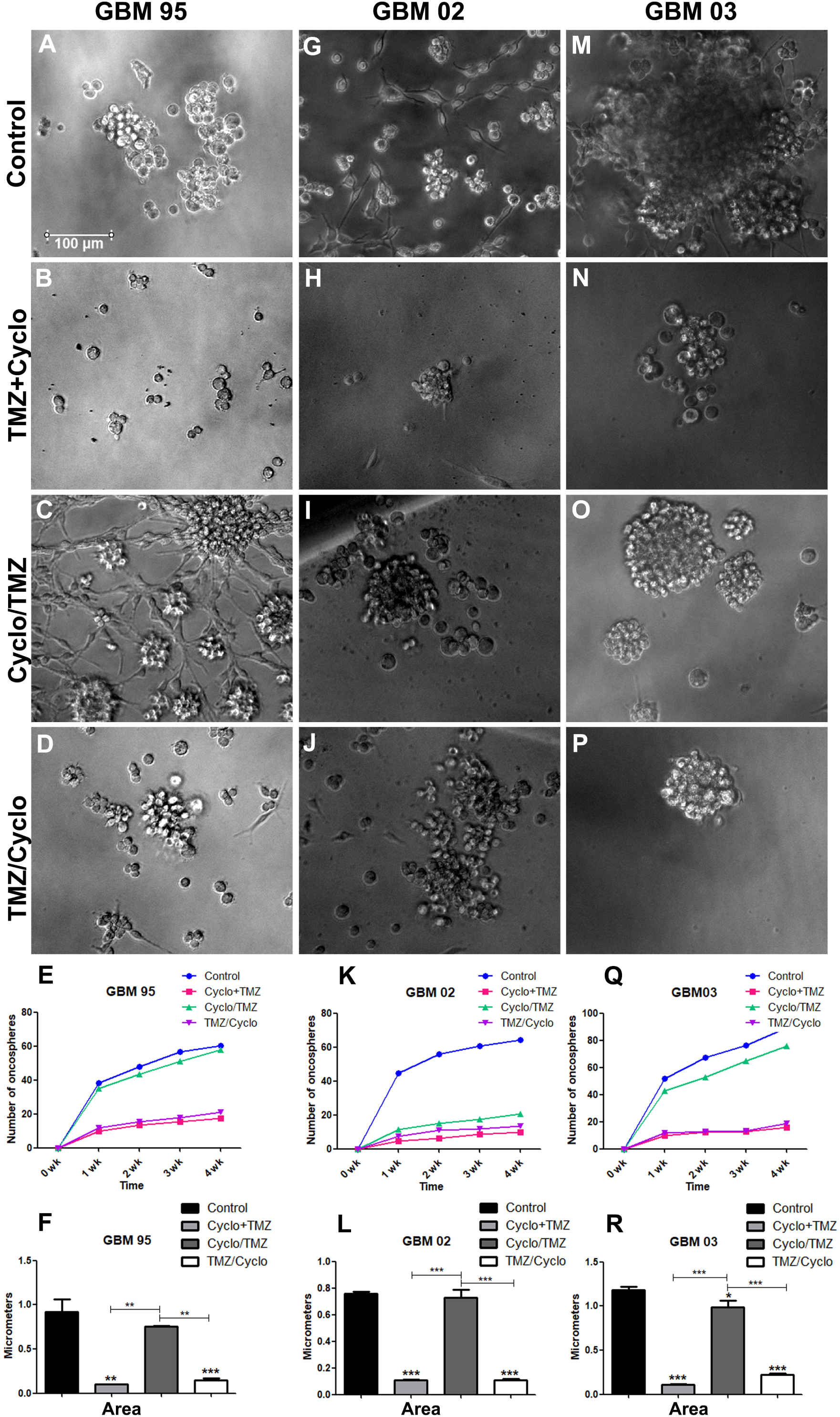
The GBM03 cell line has higher clonogenic potential. The clonogenicity assay was performed in GBM cells submitted to different combinations of cyclopamine and TMZ. The total number of colonies formed was observed using phase-contrast microscopy (**A** to **P**). The total number of formed colonies in all GBM cell lines was counted for 4 weeks (**E, K and, R**). All conditions in the three cell lines formed clones; however, the ‘exchange condition’ cells treated first with cyclopamine presented a higher number of clones (**E, K**, and **Q**). **F, L**, and **R** represent the average size of oncospheres formed by using ImageJ. GBM03 presented larger oncospheres as compared to GBM95 and GBM02 oncospheres (**F, L**, and **R**). As compared to the control conditions of the three cell lines, the GBM03 cell line has a higher number of clones than GBM02 and GBM95 (**E, K**, and **Q**). All the mean values were derived from three independent experiments. (*P< 0.05; **P< 0.01 and ***P< 0.001).

Indeed, only GBM95-derived oncospheres presented an increase of the nuclear expression of OCT-4 in the ‘exchange condition’ where TMZ was first applied (TMZ/Cyclo) (**Fig 6 D and G and Supplementary Table 5**). Although all GBM-derived oncospheres presented increased expression of SOX-2 and OCT-4 after the combination treatment (TMZ+Cyclo), instead of the cells first treated with cyclopamine and than with TMZ (Cyclo/TMZ) their expression was reduced (**Fig 6**).

**Figure 6:**
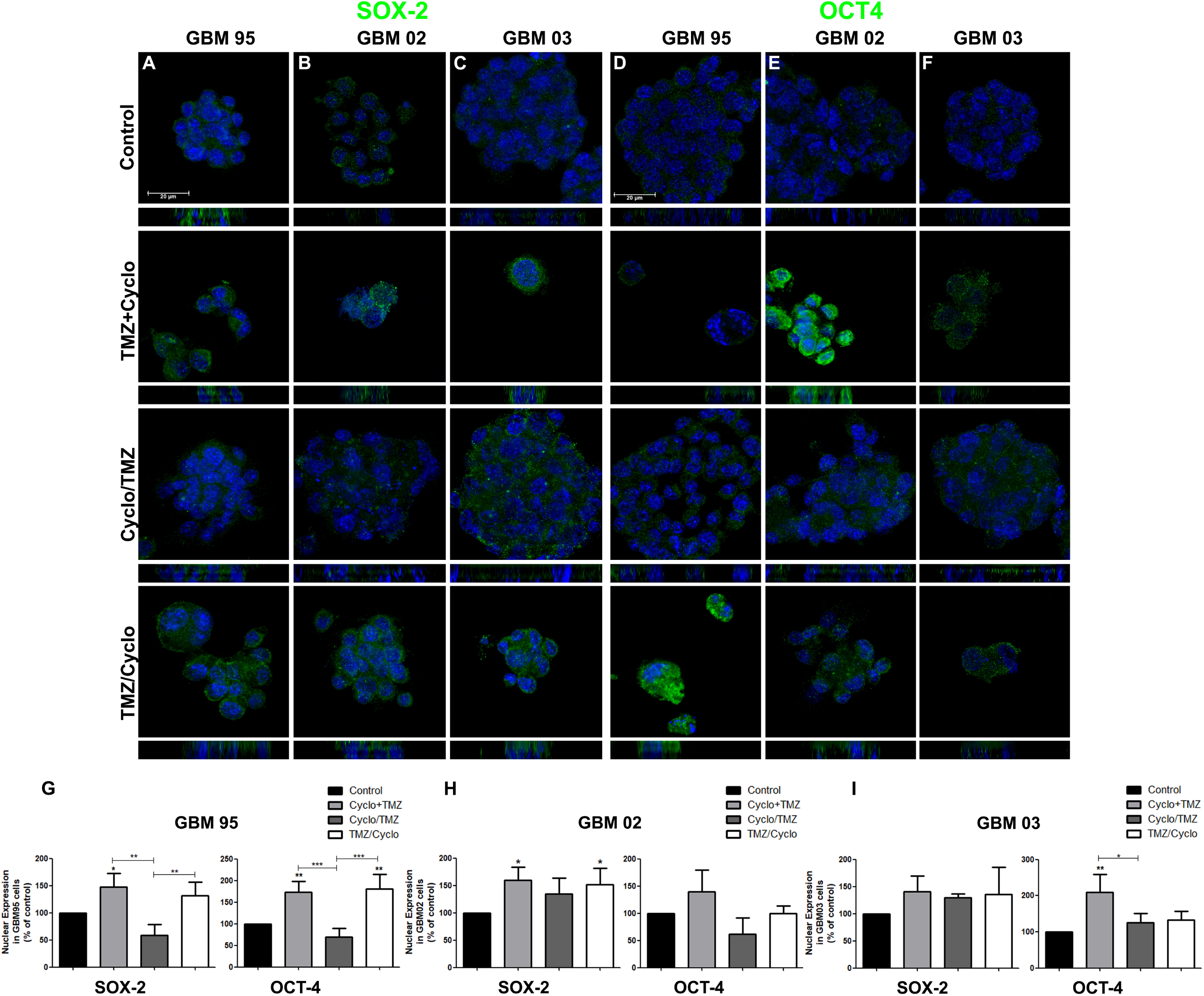
The GSCs isolated from GBM treated with cyclopamine followed by TMZ present a decrease of SOX-2 and OCT-4 nuclear expression. The expressions of SOX-2 (green) and OCT-4 (green) were observed in GSCs isolated from GBM and treated with combinations of cyclopamine and TMZ by immunofluorescence. The GBM treated cells were cultured in a defined NS34 medium and GSCs were formed after 2 weeks. Groups **A** and **D** represent nuclear expressions of SOX-2 and OCT-4 for GBM95 cell line, group **B** and **E** for GBM02 cell line, and group **C** and **F** for GBM03 cell line. All GSCs are derived from GBM cell lines treated in the ‘exchange condition’ condition, which means GBM cells treated with cyclopamine followed by TMZ treatment (Cyclo/TMZ). We observed a formation of GSCs with decreased levels of SOX-2 and OCT-4 in nuclei (**A** to **F**). The analysis of immunofluorescence of SOX-2 and for OCT4 for GBM cell lines are represented in **G, H, I** as compared to the control conditions. The cell nuclei were stained with DAPI (blue). All the mean values were derived from four independent experiments (*P< 0.05; **P< 0.01 and ***P< 0.001). Scale bar corresponds to 20 µm.

To better characterize the GSCs, we performed a differentiation assay [5,37]. The GSCs were induced to a glial differentiation by replacing the NS34 with DMEM/F-12 supplemented with 2% SFB for 10 days. After that, we analyzed the expression of GFAP, which is a marker of glial cells [38]. The cells presented typical morphological characteristics as glial cells by losing their properties of floating spheres, growing in monolayer and expressing GFAP **Supplemental Fig. 3**). It was already demonstrated that when GSCs were differentiated they become adherent and express GFAP [5].

## 4. Discussion

Patients with GBM are still confined merely to the ineffective TMZ treatment. Probably this ineffectiveness is due to the presence of GSCs which contributes to the radio and chemoresistance [5]. Previous studies have shown that TMZ is inefficient in inducing GSC death [6,7]. Indeed, several studies already showed that a combination of conventional chemotherapeutics with other anticancer drugs should be used as a therapeutic approach for GBM [27,39,40]. However, the effect of the combination of TMZ with SHH pathway inhibitors such as cyclopamine is still poorly understood.

The use of cyclopamine as SHH pathway inhibitor *in vitro* is already established for many tumor cell lines such as bone, pancreas and brain and the range of cyclopamine concentrations usually used is 5µM and 10μM [41–44]. In this work, we used an intermediary concentration of 7.5 μM cyclopamine according to MTT assay results. Although many works treated the tumor cells, such as GBM and ovarian cancer, with cyclopamine (5µM to 10µM, respectively) or in combination with other compounds for 24, 48, 72, and 96 hours [41,42,44,45]. However, there is a lack of information about what happened after 96 hours on those cells. Here, we intended to perform a cyclopamine treatment in GBM cells *in vitro* by using a similar incubation period of chemotherapy (chemotherapy protocol for GBM patients lasts 5 days) as it is applied in the clinic for GBM diagnosed patients (Brazilian National Institute of Cancer - INCA). In our study, the reduction of GBM cell viability observed after cyclopamine treatment for 192 hours (8 days) do not corroborate with the findings from Braun and coworkers, where they demonstrated that the toxicity of cyclopamine toward GBM cells is independent Hh signaling [45]. In fact, we believe that the difference observed is due to the treatment conditions used by them, since they used 24 hours of incubation period compared to our treatment condition of 192 hours (8 days).

It is interesting to note that cyclopamine had no effect on the human foreskin fibroblast cell line, suggesting that this inhibitor is not toxic for non-tumor cells, even if they have a high proliferation rate as cancer cells. This result is in accordance with another study that showed the non-toxic effect of cyclopamine in human astrocytes [46]. Demonstrating that cyclopamine has a specific effect on cancer cells instead of healthy cells [47].

GBMs patients usually present tumor recurrence, mainly during and after chemotherapy [48–50]. The mechanisms that regulate tumor recurrence are far from being understood. However, the presence of CSCs has been associated with tumor recurrence [51]. Our findings showed that cyclopamine has a prolonged effect in GBMs cell lines in ‘long term MTT’. The cell viability of all GBMs did not increase, suggesting that the cyclopamine can have a sustained effect in our GBM cell lines. Moreover, the three GBMs cell lines isolated from different patients used in this study present a pool of GSCs. We believe that the presence of GSCs could explain the slower proliferation rate observed after cyclopamine withdrawal since those cells can be “rigid”, which means they are more invasive, dividing asymmetrically and quiescent [52–54]. Another mechanism has been associated with the proliferation rate in cancer cells is TP53 mutation. In fact, GBMs cell lines used in this study, present the mutational status of TP53, which can be correlated with the more stem-like phenotype [55].

Considering that none of the patients for whom we obtained tumor tissue to establish the GBM cell lines in our laboratory were submitted to chemotherapy with TMZ, we verified if the TMZ was capable of reducing the GBM cell viability. As expected, a significant viability reduction in all GBM cell lines treated with 200μM TMZ for 144 hours was observed (6 days-time relatively similar to chemotherapy treatment)[56]. Previous reports have demonstrated the effectiveness of the combination of higher concentrations of TMZ with other compounds, such as cyclopamine in GBM cells [25,57]. However, the need to use lower concentrations of TMZ associated with other compounds to better reduce the side effects as compared to TMZ monotherapy has been exploited [27,58,59]. Recently, we have demonstrated that the selective inhibition of the SHH pathway with 20μM GANT-61 in GBM cell lines, increase the oxidative stress damage potentiating TMZ effect (400μM) and inducing cell death [60].

Herein, we compared the effect of the GBM cells treated with a combination of TMZ and cyclopamine at the same time (TMZ+Cyclo) with two other ‘exchange conditions’. Thus, in ‘exchange condition’, the GBM cells were first submitted to cyclopamine followed by TMZ treatment (Cyclo/TMZ) or vice-versa (TMZ/Cyclo). In all combined treatment conditions, we observed a significant decrease in cell viability, as described in previous studies [25,42]. The pharmacological combination of TMZ and cyclopamine (TMZ+Cyclo) showed a cooperative effect, corroborating the idea that the pharmacological combination of TMZ with other drugs, such as SHH inhibitor GANT61, are more successful than monotherapy by decreasing the cell viability of GBM cells [1,60–62].

Regarding the importance of the SHH signaling pathway for the maintenance of stemness state in tumor cells by regulating the activation of transcription factors, such as SOX-2 and OCT-4. Taking into account the presence of CSCs in GBM cell lines used, we hypothesized whether the expression of those proteins could be downregulated in GBM cells submitted to the combination of cyclopamine and TMZ [11,14,63–65]. The expressions of those proteins were distinct in GBM-treated cells after combination treatment with cyclopamine and TMZ. As we expected, GBM95 and GBM03 expressed more SOX-2 and OCT-4 than GBM02 in different treatment conditions, which could be explained by the origin of those cell lines, since they were obtained from a recurrent tumor, being the cells selected through surgery and radiotherapy. Probably, the reduced levels of SOX-2 and OCT-4 observed in GBM02 could be explained by their origin since they derived from secondary GBM that usually are less aggressive since the evolution of the disease takes longer. The secondary GBMs have different mutations compared to a primary GBM which are usually TP53 and IDH1 mutations and 19q loss [66,67]. Actually, GBM02 has IDH1 wild type, whereas GBM95 and GBM03 have IDH1 mutated, confirming our hypothesis according to the origin of the cells.

Regarding the presence of a subpopulation of GBM cells non-sensitive to the treatments (CSCs), we raised the hypothesis that other signaling pathways important for tumorigenesis could be modulating the expression of SOX-2 and OCT-4. Previous works demonstrated that WNT/β-catenin, TGF-β, epidermal growth factor receptor (EGFR) pathways could crosstalk with SHH signaling pathway, which means that depending of the tumor more than one of those signaling pathways could be activated [11,18,68,69]. It is also known that WNT/β-catenin could interact with SHH pathway through GLI1 and GLI2 by positively regulating the expression of the secreted frizzled-related protein (sFRP-1) and through GSK3β (a component of the WNT signaling pathway) inhibiting SHH signaling [11,70]. On the other hand, in some tumors when the SHH pathway is inhibited the upregulation of the WNT signaling pathway occurs in turn [71]; however, further studies will be necessary to address this hypothesis. It is possible that the increased expression of PTCH1 and SHH observed in ‘exchange condition’ may have occurred due to the interaction with other signaling pathways such as WNT and TGF-β, which may be maintaining the SHH pathway active even after treatment. In the future, more experiments should be performed to explore in detail this hypothesis.

In GBM cells submitted to ‘exchange condition’, we observed a higher number of clones in GBM-treated cells with cyclopamine followed by TMZ. However, unexpectedly, results showed that those GBM-treated cells presented decreased nuclear expression of SOX-2 and OCT-4. Thus, we suggested that the inhibition of the SHH pathway followed by TMZ administration would be a better treatment approach for GBM patients. Here, we propose a new first-line treatment with cyclopamine by acting on SMO and consequently causing the inhibition of the SHH pathway followed by TMZ treatment. We believe that this therapeutic approach will reduce the aggressiveness of the tumor cells by sensitizing the GSCs for TMZ treatment. Ultimately, the combination of cyclopamine with TMZ treatment at the same time induces the expression of SOX-2 and OCT-4 in GBM cells, suggesting that this approach leads to the stemness and resistance of GBM cells to treatments and may result in tumor relapse in patients. Herein, we suggest that the best therapeutic strategy would be to inhibit the SHH pathway first, followed by TMZ administration. Nevertheless, in the future, efforts should be made to investigate the possibility of a therapeutic approach using SHH inhibitors before conventional chemotherapy in GBM patients.

## Abbreviations

BSA: bovine serum albumins;
CDI: coefficient of drug interaction;
DAPI: 4-6-diamino-2-phenylindole;
DMEM/F12: Dulbecco’s Modified Eagle Medium/Nutrient Mixture F-12;
DMSO: dimethyl sulfoxide;
EGFR: epidermal growth factor receptor;
FBS: fetal bovine serum;
GBM: glioblastoma;
GFAP: glial fibrillary acidic protein;
GSCs: glioma stem cells;
INCA: National Institute of Cancer;
MGMT: O-6-methylguanine-DNA methyltransferase;
MTT: 3-(4, 5-dimethylthiazol-2-yl)-2,5-diphenyl tetrazolium bromide;
PFA: paraformaldehyde;
PTCH: Patched;
PVDF: hybond-P polyvinylidene difluoride;
sFRP-1: secreted frizzled related protein;
SHH: Sonic hedgehog;
SMO: Smoothened;
TBS-T: tris-buffered saline with tween;
TGF-β: transforming growth beta;
TME: tumor microenvironment;
TMZ: temozolomide;
WNT: wingless.

## Author contributions statements

The manuscript was conceptualized, written, edited and critically evaluated by each of the authors. GC performed all experiments. JR contributed to *western-blotting*. LP contributed to the qRT-PCR. AN contributed to the search for clinical patients’ history and article final preparation. The work was supervised by TS and DM. All authors read and approved the final submitted version of the manuscript.

## Acknowledgments

We thank Professor Vivaldo Moura Neto for carefully reading the manuscript and all the support given. This work was supported by the National Institute for Translational Neuroscience (INNT) of the Ministry of Science and Technology; Brazilian Federal Agency for the Support and Evaluation of Graduate Education (CAPES) of the Ministry of Education; National Council for Scientific and Technological Development (CNPq); Fundação Carlos Chagas Filho de Amparo à Pesquisa do Estado do Rio de Janeiro Carlos Chagas Filho Research Support Foundation (FAPERJ); AryFrauzino Foundation for Cancer Research and Pró-SaúdeAssociaçãoBeneficente de Assistência Social e Hospitalar

## Supplementary data

**Supplementary Figure 1**: ***GBM cell viability after TMZ treatments by MTT assay*.** The cell viability was analyzed after the TMZ treatment at different concentrations of TMZ (100 µM, 200 µM, 400 µM, and 600 µM) in GBM95, GBM02, and GBM03 for 72 hours (**A**) and in GBM95, GBM02, and GBM03 for 144 hours (**B**) (6 days). After 72 hours of TMZ treatment, only GBM 02 and GBM03 showed a decrease in cell viability (**A**). After 144 hours of treatment, all three cell lines showed a significant decrease of cell viability for higher concentrations of 200 µM (**B**). The average number of treatment-resistant cells to cyclopamine (5 μM, 7.5 μM, and 10μM) was assessed in GBM95, GBM02, and GBM03 after 192 hours (**C**) (8 days) of treatment counting the positive cells for DAPI through fluorescence microscopy. The average number of treatment-resistant cells to cyclopamine and TMZ after 144 hours on GBM95, GBM02, and GBM03 are represented in (**D**) where we counted the positive cells for DAPI through fluorescence microscopy stained with DAPI. All the mean values are derived from three independent experiments after the MTT assays. The decrease in cell viability was analyzed comparing all conditions with the control condition. (*P< 0.05; **P< 0.01 and ***P< 0.001).

**Supplementary Figure 2: *GBM03 cell line presented more GSCs*.** The capability of forming floating oncospheres in GBM cell lines after treatment with different combinations of cyclopamine with TMZ was observed by using phase-contrast microscopy. **A** to **D** represent GBM95 cell line, **F** to **I** the GBM02 cell line, and **K** to **N** the GBM03 cell line. All cell lines formed oncospheres after two weeks in culture with NS34 medium; however, the GBM03 cell line formed a higher amount of GSC in control condition (**K** and **O**), when compared to the control conditions of GBM95 and GBM02 (**A, E**, and **F, J**, respectively). For all GBMs, there was a decrease in floating oncospheres in different combination treatments of cyclopamine with TMZ when compared with the control condition. All the mean values were derived from three independent experiments (*P<0.05; **P<0.01 and ***P<0.001). Scale bar corresponds to 50 µm.

**Supplementary Figure 3: *GFAP expression in GSCs derived from GBM treated cells after differentiation assay*.** GFAP expression in differentiated cells from GSCs of GBM treated cells was assessed by immunofluorescence. The GBM treated cells were cultured in a defined NS34 medium, and GSCs were formed after 2 weeks. The GSCs were then cultured in DMEM with 2% FBS to differentiate the cells in astroglial cells. After the differentiation assay, all cells are GFAP-positive (GBM95, **A-D**; GBM02 **E-H**; and GBM03 **I-L**). The cell nuclei were stained with DAPI (blue). Scale bar corresponds to 30 µm.

## Tables

**Table 1-**Synergistic effects of TMZ and Cyclopmine in human GBM cells

**Supplementary Table 1**-Effect of Cyclopamine GBM cells (cell viability)

**Supplementary Table 2**-Effect of Cyclopamine GBM cells (cell viability – long term MTT)

**Supplementary Table 3**-Effect of Cyclopamine and TMZ on GBM cells (cell viability)

**Supplementary Table 4**-Effect of Cyclopamine and TMZ on GBM cells

**Supplementary Table 5**-Effect of Cyclopamine and TMZ on GBM cells (oncospheres)

**Supplementary Table 6**-Effect of TMZ on GBM cells

