## Supplementary figures and images for "Cyclopamine sensitizes Glioblastoma cells to Temozolomide treatment through Sonic Hedgehog pathway"

### Supplemental figure 1

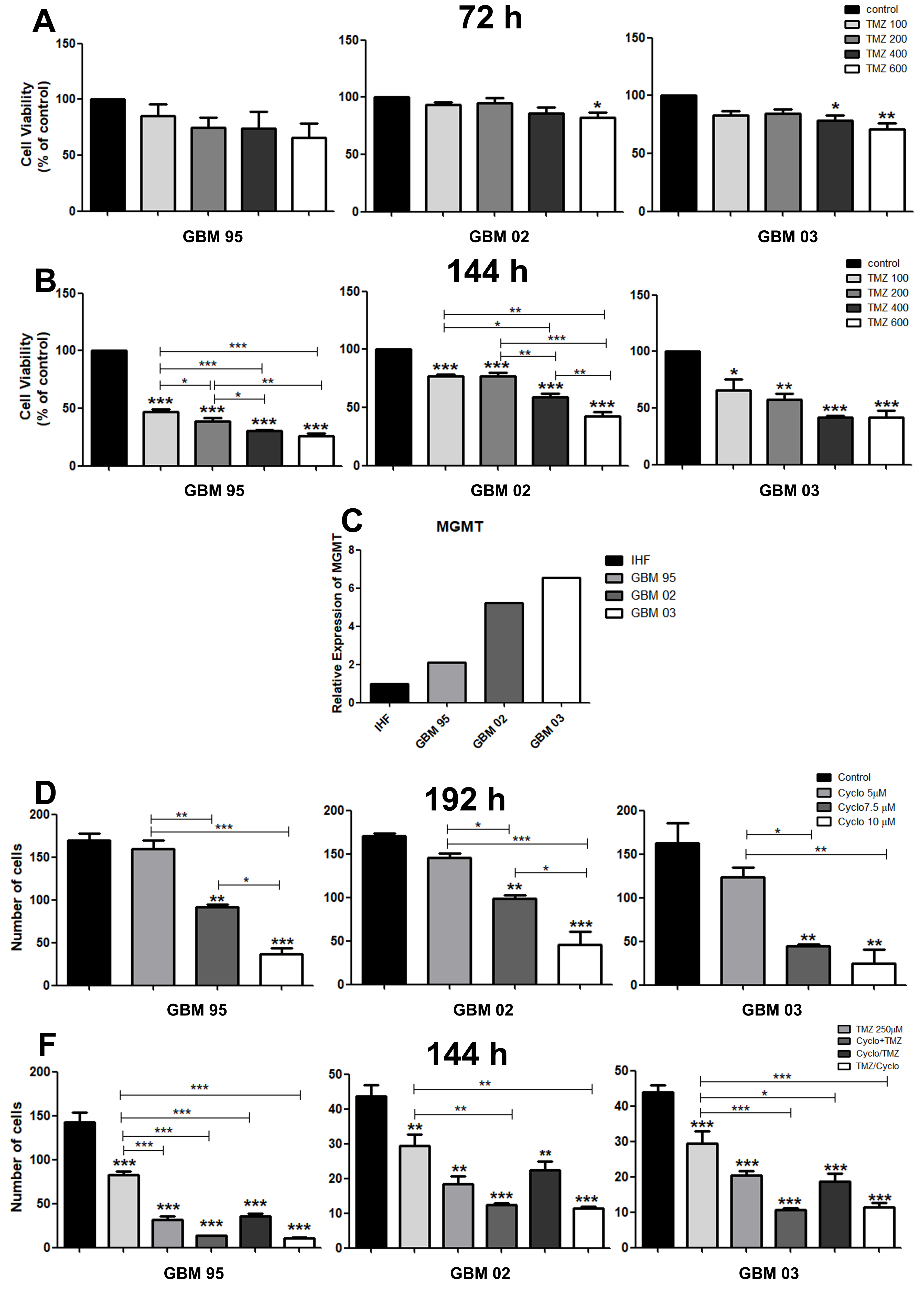

### Supplemental Figure 2

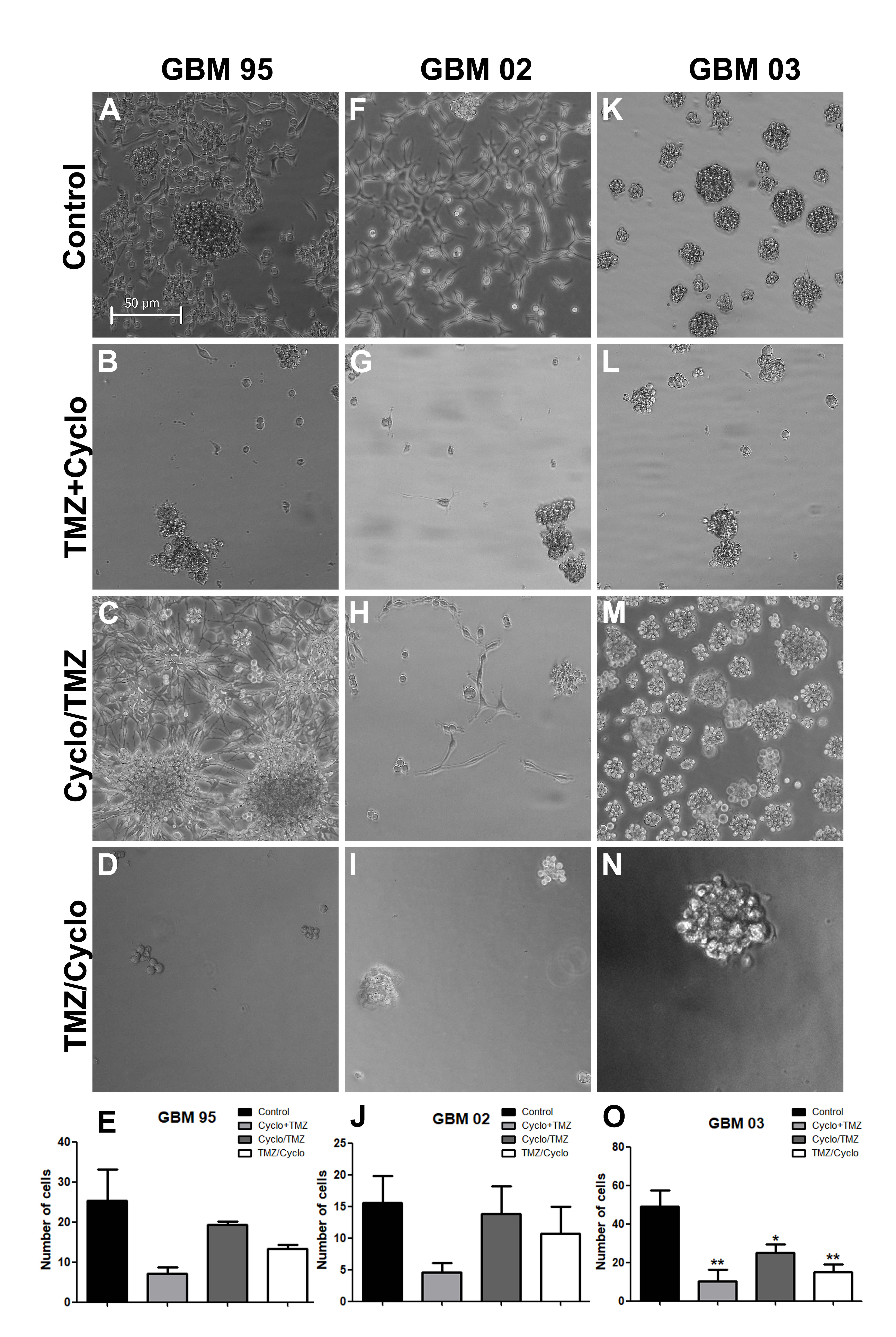

### Supplemental Figure 3

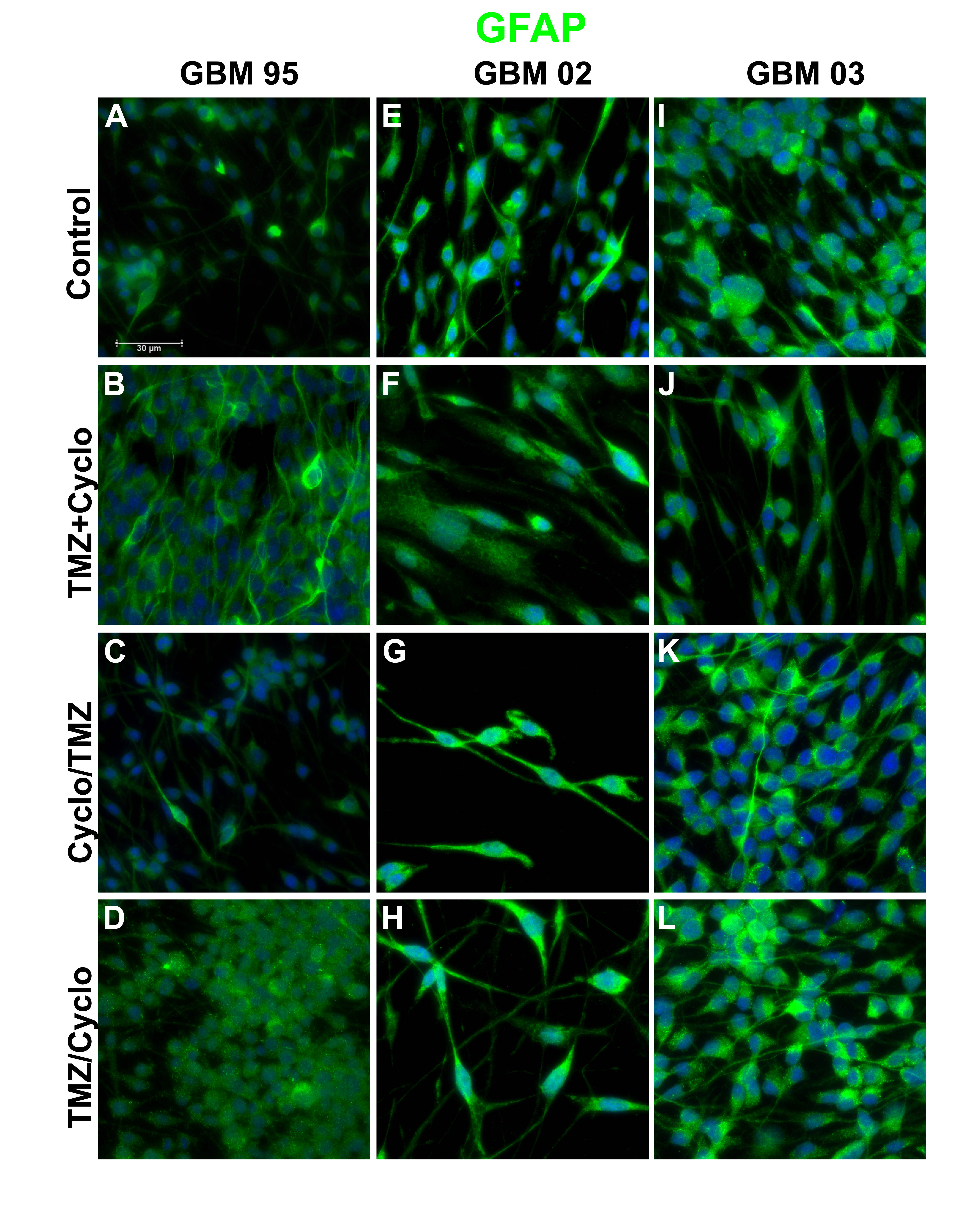
