## Supplemental Tables for "Cyclopamine sensitizes Glioblastoma cells to Temozolomide treatment through Sonic Hedgehog pathway"

**Supplementary Table 1-** Effect of Cyclopamine GBM cells (cell viability)

|  | **GBM 95** | | | | **GBM 02** | | | | **GBM 03** | | | |
| --- | --- | --- | --- | --- | --- | --- | --- | --- | --- | --- | --- | --- |
|  | *48h* | *96h* | *192h* | *Number of cells (192h)* | *48h* | *96h* | *192h* | *Number of cells (192h)* | *48h* | *96h* | *192h* | *Number of cells (192h)* |
| *Control vs 5μM* | 20.71±1.49 | 17.08±1.22 | 27.21±1.95 | 9.97±1.30 | 41.74±3.37* | 7.84±0.63 | 4.00±0.32 | 25.23±2.94 | 20.22±1.41 | 2.58±0.18 | 18.56±1.30 | 38.31±2.52 |
| *Control vs 7.5μM* | 57.63±4.14** | 50.17±3.60** | 41.49±2.98* | 78.65±9.23** | 46.30±3.74 | 17.11±1.38 | 8.78±0.71 | 71.96±7.50** | 63.68±4.46*** | 39.09±2.74* | 69.35±4.86*** | 117.5±7.74** |
| *Control vs 10μM* | 65.78±4.73*** | 71.36±5.13*** | 69.15±4.97*** | 133.1±17.46*** | 2.65±0.21 | 1.88±0.15 | 40.56±3.28 | 125.4±14.62*** | 73.13±5.12*** | 22.68±1.59 | 80.30±5.63*** | 137.8±9.08*** |
| *5μM vs 7.5μM* | 36.93±2.65* | 33.09±2.38 | 14.28±1.02 | 68.68±8.06** | 12.98±0.25 | 24.53±0.48 | 33.05±0.65 | 46.73±4.87* | 43.46±3.04* | 36.50±2.55 | 50.79±3.56** | 79.21±5.22* |
| *5μM vs 10μM* | 45.07±3.2* | 54.28±3.9** | 41.94±3.01* | 123.1±16.15*** | 151.0±3.0* | 62.48±1.24 | 49.16±0.98 | 100.2±11.68*** | 52.90±3.70** | 20.09±1.40 | 61.74±4.32*** | 99.48±6.55** |
| *7.5μM vs 10μM* | 8.14±0.58 | 21.19±1.52 | 27.66±1.99 | 54.40±6.38* | 138.0±2.75* | 37.96±0.75 | 16.11±0.32 | 53.43±5.57* | 9.44±0.66 | 16.41±1.15 | 10.95±0.76 | 20.27±1.33 |

*P< 0.05; **P< 0.01 and ***P< 0.001

**Supplementary Table 2-** Effect of Cyclopamine GBM cells (cell viability – long term MTT)

|  | **GBM 95** | | | **GBM 02** | | | **GBM 03** | | |
| --- | --- | --- | --- | --- | --- | --- | --- | --- | --- |
|  | *8 day* | *12 day* | *16 day* | *8 day* | *12 day* | *16 day* | *8 day* | *12 day* | *16 day* |
| *Control vs 5μM* | 43.34±2.59* | 56.40±3.37** | 55.81±3.34** | 47.57±2.87* | 31.97±1.93 | 9.90±0.59 | 35.28±2.47* | 56.08±3.93*** | 50.51±3.54** |
| *Control vs 5μM (w/o)* | 43.34±2.59* | 56.02±3.35** | 60.19±3.60** | 47.57±2.87* | 26.08±1.57 | 4.05±0.24 | 35.28±2.47* | 39.08±2.74* | 31.14±2.18 |
| *Control vs 7.5μM* | 64.46±3.85*** | 81.73±4.89*** | 71.89±4.30*** | 61.20±3.69** | 79.19±4.36*** | 46.44±2.80* | 45.83±3.21** | 60.35±4.23*** | 71.98±5.05*** |
| *Control vs 7.5μM*  *(w/o)* | 64.46±3.85*** | 76.86±4.60*** | 63.97±3.82*** | 61.20±3.69** | 62.71±3.78*** | 19.21±1.16 | 45.83±3.21** | 58.48±4.10*** | 66.50±4.66*** |
| *Control vs 10μM* | 75.51±4.52*** | 93.79±5.61*** | 92.13±5.51*** | 71.51±4.32*** | 89.39±5.40*** | 88.15±5.32*** | 46.38±3.25** | 75.14±5.27*** | 86.64±6.07*** |
| *Control vs 10μM*  *(w/o)* | 75.29±4.50*** | 91.20±5.45*** | 82.29±4.92*** | 71.51±4.32*** | 76.80±4.64*** | 61.72±3.72** | 46.38±3.25** | 69.28±4.86*** | 80.50±5.64*** |
| *5μM vs 5μM*  *(w/o)* | - | 0.38±0.02 | 4.38±0.26 | - | 5.88±0.35 | 5.84±0.35 | - | 17.00±1.19 | 19.37±1.35 |
| *5μM vs 7.5μM* | 21.11±1.26 | 25.33±1.51 | 16.08±0.96 | 13.63±0.82 | 40.23±2.43 | 36.53±2.20 | 10.56±0.74 | 4.27±0.29 | 21.47±1.50 |
| *5μM vs 7.5μM*  *(w/o)* | 21.11±1.26 | 20.46±1.22 | 8.16±0.488 | 13.63±.0.82 | 30.74±1.85 | 9.30±0.56 | 10.56±0.74 | 2.40±0.16 | 15.99±1.12 |
| *5μM vs 10μM* | 32.17±1.92 | 37.39±2.23 | 36.32±2.17 | 23.94±1.44 | 57.43±3.47** | 78.24±4.72*** | 11.10±0.77 | 19.05±1.33 | 36.13±2.53* |
| *5μM vs 10μM*  *(w/o)* | 31.94±1.91 | 34.80±2.08 | 26.49±1.58 | 23.94±1.44 | 44.83±2.70* | 51.82±3.13** | 11.10±0.77 | 13.20±0.92 | 29.99±2.10 |
| *5μM (w/o) vs 7.5μM* | 21.11±1.26 | 25.71±1.53 | 11.70±0.70 | 13.63±0.82 | 46.12±2.78* | 42.38±2.56* | 10.56±0.74 | 21.27±1.49 | 40.83±2.86* |
| *5μM (w/o) vs 7.5μM (w/o)* | 21.11±1.26 | 20.84±1.14 | 3.78±0.22 | 13.63±0.82 | 36.63±2.21 | 15.16±0.91 | 10.56±0.74 | 19.40±1.36 | 35.36±2.48* |
| *5μM (w/o) vs 10μM* | 32.17±1.92 | 37.77±2.26 | 31.94±1.91 | 23.94±1.44 | 63.31±3.82*** | 84.09±5.08*** | 11.10±0.77 | 36.06±2.53* | 55.50±3.89** |
| *5μM (w/o) vs 10μM (w/o)* | 31.94±1.91 | 35.19±2.10 | 22.10±1.32 | 23.94±1.44 | 50.72±3.06** | 57.67±3.48** | 11.10±0.77 | 30.20±2.11 | 49.36±3.46** |
| *7.5μM vs 7.5μM*  *(w/o)* | - | 4.8±0.29 | 7.91±0.47 | - | 9.48±0.57 | 27.22±1.64 | - | 1.86±0.13 | 5.47±0.38 |
| *7.5μM vs 10μM* | 11.05±0.66 | 12.06±0.72 | 20.24±1.21 | 10.31±0.62 | 17.20±1.03 | 41.71±2.52* | 0.54±0.03 | 14.79±1.03 | 14.66±1.02 |
| *7.5μM vs 10μM*  *(w/o)* | 10.83±0.64 | 9.47±0.56 | 10.41±0.62 | 10.31±0.62 | 4.60±0.27 | 15.28±0.92 | 0.54±0.03 | 8.93±0.62 | 8.52±0.59 |
| *7.5μM (w/o) vs 10μM* | 11.05±0.66 | 16.93±1.01 | 28.16±1.68 | 10.31±0.62 | 26.69±1.61 | 68.93±4.16*** | 0.54±0.03 | 16.65±1.16 | 20.14±1.41 |
| *7.5μM (w/o) vs 10μM (w/o)* | 10.83±0.64 | 14.34±0.85 | 18.32±1.09 | 10.31±0.62 | 14.09±0.85 | 42.51±2.56* | 0.54±0.03 | 10.80±0.75 | 14.00±0.98 |
| *10μM vs 10μM*  *(w/o)* | 0.22±0.01 | 2.58±0.15 | 9.83±058 | - | 12.59±0.76 | 26.43±1.59 | - | 5.85±0.41 | 6.13±0.43 |

*P< 0.05; **P< 0.01 and ***P< 0.001

**Supplementary Table 3-** Effect of Cyclopamine and TMZ on GBM cells (cell viability)

|  | **GBM 95** | | | **GBM 02** | | | **GBM 03** | | |
| --- | --- | --- | --- | --- | --- | --- | --- | --- | --- |
|  | *48h* | *72h* | *144h* | *48h* | *72h* | *144h* | *48h* | *72h* | *144h* |
| *Control vs Cyclo 7.5 µM* | 20.81±3.58** | 20.04±3.45** | 33.76±5.82*** | 15.87±1.50 | 28.25±2.67* | 34.28±3.24* | 13.08±1.22 | 28.82±2.69* | 69.77±6.53** |
| *Control vs TMZ 250µM* | 31.29±5.39*** | 40.96±7.06*** | 61.38±10.58*** | 35.86±3.39** | 38.89±3.68** | 23.34±2.11 | 21.63±2.02 | 32.49±3.04* | 42.78±4.00** |
| *Control vs Cyclo+TMZ* | 12.23±2.11 | 35.05±6.04*** | 77.30±13.33*** | 10.43±0.98 | 19.12±1.81 | 80.93±7.66*** | 10.09±0.94 | 21.95±2.05 | 77.56±7.26*** |
| *Cyclo 7.5 µM vs TMZ 250µM* | 10.48±1.80 | 20.92±3.60** | 27.62±4.76*** | 19.98±1.89 | 10.64±1.00 | 10.94±1.03 | 8.54±0.80 | 3.67±0.34 | 26.99±2.52 |
| *Cyclo 7.5 µM vs Cyclo+TMZ* | 8.57±1.47 | 15.01±2.58* | 43.54±7.50*** | 5.44±0.51 | 9.13±0.86 | 46.65±4.41*** | 2.99±0.28 | 6.87±0.64 | 7.79±0.72 |
| *TMZ 250µM vs Cyclo+TMZ* | 19.06±3.28* | 5.91±1.01 | 1592±2.74* | 25.42±2.40 | 19.77±1.87 | 57.59±5.45*** | 11.53±1.08 | 10.54±0.98 | 34.78±3.25* |
| *Control vs Cyclo/TMZ* | 26.23±4.16** | 38.80±6.16*** | - | 18.88±2.75* | 26.56±3.88** | - | 24.63±5.98*** | 46.01±11.18*** | - |
| *Control vs TMZ/Cyclo* | 35.50±5.64*** | 62.37±9.91*** | - | 20.41±2.98* | 28.42±4.15** | - | 37.25±9.05*** | 55.25±13.42*** | - |
| *Cyclo/TMZ vs TMZ/Cyclo* | 9.26±1.47 | 23.58±3.74** | - | 1.52±0.22 | 1.85±0.27 | - | 12.62±3.06* | 9.24±2.24 | - |

*P< 0.05; **P< 0.01 and ***P< 0.001

**Supplementary Table 4-** Effect of Cyclopamine and TMZ on GBM cells

|  | **GBM 95** | | | | | | | | **GBM02** | | | | | |
| --- | --- | --- | --- | --- | --- | --- | --- | --- | --- | --- | --- | --- | --- | --- |
|  | *SOX-2 % (IHQ)* | *OCT-4 %*  *(IHQ)* | *SOX-2 % (WB)* | *OCT-4 %*  *(WB)* | *PTCH1 % (WB)* | *SHH %*  *(WB)* | *Casp 3 %*  *(WB)* | *Number of cells* | *SOX-2 % (IHQ)* | *OCT-4 %*  *(IHQ)* | *SOX-2 % (WB)* | *OCT-4 %*  *(WB)* | *PTCH1 % (WB)* | *SHH %*  *(WB)* |
| *Control vs Cyclo 7.5 µM* | 86.4±4.7 | 31.6±2.6 | 27.8±2.1 | 18.6±1.6 | 35.36±1.8 | 88.7±3.5 | 4.8±0.9 | 60.1±10.2 *** | 1.7±0.1 | 28.2±1.6 | 36.7±4.0 | 62.9±4.5 | 9.2±0.5 | 24.6±1.0 |
| *Control vs TMZ 250µM* | 35.5±1.9 | 1.9±0.1 | 99.2±7.7  ** | 67.5±5.7  * | 38.4±2.0 | 34.5±1.3 | 7.3±1.4 | 110.8±17.9 *** | 154.1±9.2 *** | 142.4±8.0 ** | 43.6±4.2 | 20.4±1.6 | 50.1±2.5 | 17.1±0.7 |
| *Control vs Cyclo+TMZ* | 132.1±7.2  ** | 47.1±3.9 | 99.9±7.5  ** | 33.3±2.5 | 59.8±3.1 | 152.1±5.4* | 14.7±2.9 | 129.4±20.9 *** | 130.8±7.8 ** | 137.6±7.7 ** | 8.3±0.8 | 105.6±7.5  ** | 68.4±3.5 | 63.3±2.6 |
| *Control vs Cyclo/TMZ* | 18.9±1.1 | 119.2±8.6  *** | 88.2±6.6  ** | 132.9±10.0  *** | 13.0±0.6 | 626.5±22.4*** | 27.4±5.4 | 107.5±17.4 *** | 58.21±4.0 | 88.7±4.4 | 14.3±1.4 | 30.9±2.2 | 5.9±0.3 | 83.4±3.5 |
| *Control vs TMZ/Cyclo* | 278.1±15.3*** | 16.4±1.3 | 160.3±10.6 *** | 106.1±8.0  ** | 61.7±3.2 | 338.6±12.1*** | 1.8±0.35 | 132.3±22.5 *** | 246.1±14.8 *** | 264.6±13.0 *** | 83.6±8.1  ** | 129.9±10.3 *** | 27.8±1.4 | 148.5±5.5  * |
| *Cyclo 7.5 µM vs TMZ 250µM* | 50.9±2.5 | 29.6±2.1 | 71.3±5.0  * | 48.7±4.1 | 3.1±0.1 | 54.1±2.2 | 12.2±2.4 | 50.7±7.9 *** | 152.5±7.6 ** | 114.2±5.7  * | 7.0±0.7 | 42.5±3.0 | 59.3±3.0 | 7.4±0.3 |
| *Cyclo 7.5 µM vs Cyclo+TMZ* | 45.6±2.2 | 15.5±1.1 | 72.1±5.1  * | 14.6±1.1 | 24.4±1.2 | 63.9±2.3 | 19.6±3.9 | 69.4±10.8 *** | 129.0±6.5 ** | 109.3±5.5  * | 28.3±2.7 | 42.7±2.8 | 77.7±3.9 | 38.7±1.6 |
| *Cyclo 7.5 µM vs Cyclo/TMZ* | 67.5±3.7 | 87.5±5.7  * | 60.3±4.2 | 114.3±8.6  ** | 48.3±2.5 | 537.8±19.2*** | 32.9±6.4* | 47.4±7.4 *** | 56.5±3.1 | 60.4±2.7 | 22.4±2.2 | 32.0±2.0 | 3.3±0.2 | 58.8±2.4 |
| *Cyclo 7.5 µM vs TMZ/Cyclo* | 191.7±9.6  ** | 15.2±1.1 | 132.5±8.3  *** | 87.4±6.6  * | 26.4±1.3 | 249.9±8.9  ** | 6.7±1.3 | 72.3±11.9 *** | 244.4±12.3*** | 236.3±10.7 *** | 46.9±4.5 | 66.9±4.7 | 18.6±0.9 | 124.0±4.6 |
| *TMZ 250µM vs Cyclo+TMZ* | 96.5±4.8 | 45.1±3.3 | 0.7±0.04 | 34.2±2.6 | 21.3±1.1 | 117.6±4.2 | 7.7±1.5 | 18.6±2.8 | 23.5±1.2 | 4.8±0.2 | 35.2±3.1 | 85.2±6.0  * | 18.3±0.9 | 46.1±1.9 |
| *TMZ 250µM vs Cyclo/TMZ* | 16.6±0.1 | 117.2±7.6  ** | 11.1±0.8 | 65.4±4.9 | 51.4±2.7 | 592.0±21.2*** | 20.1±4.0 | 3.3±0.5 | 95.9±5.3  * | 53.7±2.4 | 29.3±2.6 | 10.5±0.7 | 56.0±2.8 | 66.2±2.8 |
| *TMZ 250µM vs TMZ/Cyclo* | 242.6±12.2*** | 14.4±1.0 | 61.1±3.8 | 38.5±2.9 | 23.1±1.2 | 304.1±10.8*** | 5.5±1.1 | 21.6±3.4 | 92.0±4.6 | 122.2±5.5  * | 40.0±3.5 | 109.4±8.7  ** | 77.9±4.0 | 131.3±4.9  * |
| *Cyclo+TMZ vs Cyclo/TMZ* | 113.2±6.2  * | 72.7±4.7  * | 11.8±0.8 | 99.6±6.7  ** | 72.8±3.8 | 474.4±15.5*** | 12.9±2.5 | 21.9±3.3 | 72.5±4.0 | 48.9±2,2 | 6.0±0.5 | 74.7±4.8 | 74.3±3.8 | 20.1±0.8 |
| *Cyclo+TMZ vs TMZ/Cyclo* | 146.0±7.34  ** | 30.7±2.2 | 60.4±3.8 | 72.7±5.0  * | 1.9±0.1 | 186.4±6.1  * | 12.9±2.6 | 2.9±0.4 | 115.5±5.8  * | 127.0±5.7  * | 75.2±6.7  * | 24.15±1.7 | 96.2±4.9 | 85.2±3.2 |
| *Cyclo/TMZ vs TMZ/Cyclo* | 259.2±14.2*** | 102.8±6.7  ** | 72.2±4.5 | 26.9±1.8 | 74.7±3.9 | 287.9±9.4  *** | 25.6±5.1 | 24.9±3.9 | 187.9±10.3*** | 175.9±7.2 ** | 69.3±6.1  * | 98.9±7.0  ** | 21.9±1.1 | 65.1±2.4 |

*P< 0.05; **P< 0.01 and ***P< 0.001

**Supplementary Table 4-** Effect of Cyclopamine and TMZ on GBM cells (cont)

|  | **GBM 02** | | **GBM 03** | | | | | | | |
| --- | --- | --- | --- | --- | --- | --- | --- | --- | --- | --- |
|  | *Casp 3 %*  *(WB)* | *Number of cells* | *SOX-2 % (IHQ)* | *OCT-4 %*  *(IHQ)* | *SOX-2 % (WB)* | *OCT-4 %*  *(WB)* | *PTCH1 % (WB)* | *SHH %*  *(WB)* | *Casp 3 %*  *(WB)* | *Number of cells* |
| *Control vs Cyclo 7.5 µM* | 137.1±8.9  *** | 14.3±5.9 ** | 58.8±2.1 | 8.4±0.6 | 43.8±1.7 | 5.1±0.5 | 17.8±1.6 | 27.7±1.6 | 0.3±0.02 | 14.6±6.6 *** |
| *Control vs TMZ 250µM* | 141.4±9.2  *** | 25.2±9.5 *** | 43.9±1.6 | 20.1±1.4 | 36.5±1.4 | 5.1±0.5 | 31.2±3.1 | 57.5±3.4 | 86.2±5.9  * | 23.5±10.3 *** |
| *Control vs Cyclo+TMZ* | 147.5±9.6  *** | 31.3±12.4 *** | 255.6±9.1  ** | 43.7±3.7 | 175.7±6.8  * | 68.3±6.4  ** | 5.5±0.5 | 2.7±0.2 | 45.4±3.1 | 33.3±13.9 *** |
| *Control vs Cyclo/TMZ* | 212.8±12.4  *** | 21.2±7.9 *** | 176.4±6.3  * | 42.4±3.5 | 74.1±2.9 | 14.6±1.2 | 3.5±0.3 | 71.9±4.3 | 152.6±11.4  *** | 25.4±11.1 *** |
| *Control vs TMZ/Cyclo* | 54.1±3.1 | 32.2±11.2 *** | 112.9±4.0 | 148.9±10.9  *** | 113.9±4.4 | 81.6±7.6  ** | 27.6±2.8 | 96.9±5.7  * | 60.9±4.1 | 32.53±13.6 *** |
| *Cyclo 7.5 µM vs TMZ 250µM* | 4.3±0.3 | 11.0±3.9 | 14.9±0.5 | 11.9±0.7 | 7.3±0.3 | 0.3±0.03 | 49.2±4.4 | 29.8±1.6 | 85.9±5.8  * | 8.9±4.0 |
| *Cyclo 7.5 µM vs Cyclo+TMZ* | 10.4±0.7 | 17.0±6.4** | 196.8±6.4  * | 52.1±3.5 | 131.9±5.1 | 62.8±5.9  * | 23.4±2.1 | 25.0±1.3 | 45.0±3.1 | 18.7±8.0 *** |
| *Cyclo 7.5 µM vs Cyclo/TMZ* | 75.7±4.4 | 6.9±2.5 | 117.6±3.8 | 34.0±2.3 | 30.3±1.2 | 20.1±1.7 | 14.2±1.3 | 44.2±2.4 | 152.2±11.3  *** | 10.8±4.9 * |
| *Cyclo 7.5 µM vs TMZ/Cyclo* | 83.0±4.8 | 17.9±6.0 ** | 54.1±1.8 | 157.3±9.6  *** | 70.1±2.7 | 76.1±7.1  ** | 9.8±0.9 | 69.2±3.7 | 60.5±4.1 | 17.9±7.7 *** |
| *TMZ 250µM vs Cyclo+TMZ* | 6.0±0.4 | 6.0±2.0 | 211.8±6.9  * | 63.8±4.3 | 139.2±5.4 | 63.1±5.9  * | 25.8±2.6 | 54.8±2.9 | 40.8±2.8 | 9.8±4.0 |
| *TMZ 250µM vs Cyclo/TMZ* | 71.4±4.2 | 4.0±1.3 | 132.6±4.3 | 22.3±1.5 | 37.7±1.5 | 19.7±1.6 | 34.9±3.5 | 14.4±0.8 | 66.3±4.9 | 1.9±0.8 |
| *TMZ 250µM vs TMZ/Cyclo* | 87.3±5.1  * | 6.9±2.1 | 69.1±2.2 | 169.0±10.3  *** | 77.4±3.0 | 76.4±7.2  ** | 59.0±5.9  * | 39.3±2.1 | 25.3±1.7 | 9.0±3.8 |
| *Cyclo+TMZ vs Cyclo/TMZ* | 65.3±3.8 | 10.0±3.5 | 79.2±2.5 | 86.1±6.4  ** | 101.6±3.9 | 82.9±6.9  ** | 9.1±0.9 | 69.2±3.7 | 107.2±8.0  ** | 7.9±3.3 |
| *Cyclo+TMZ vs TMZ/Cyclo* | 93.4±5.4  * | 0.9±0.3 | 142.7±4.6 | 105.2±7.05  ** | 61.8±2.4 | 13.2±1.2 | 33.2±3.3 | 94.2±5.1 | 15.5±1.1 | 0.7±0.3 |
| *Cyclo/TMZ vs TMZ/Cyclo* | 158.7±8.5  ** | 11.0±3.4 | 63.5±2.1 | 191.4±12.9  *** | 39.8±1.5 | 96.1±8.1  ** | 24.1±2.4 | 24.9±1.3 | 91.8±6.8  * | 7.1±3.0 |

*P< 0.05; **P< 0.01 and ***P< 0.001

**Supplementary Table 5-** Effect of Cyclopamine and TMZ on GBM cells (oncosferes)

|  | **GBM 95** | | | | **GBM 02** | | | | **GBM 03** | | | |
| --- | --- | --- | --- | --- | --- | --- | --- | --- | --- | --- | --- | --- |
|  | *Area* | *SOX-2 (ICQ)* | *OCT-4 (ICQ)* | *Number of cells* | *Area* | *SOX-2 (ICQ)* | *OCT-4 (ICQ)* | *Number of cells* | *Area* | *SOX-2 (ICQ)* | *OCT-4 (ICQ)* | *Number of cells* |
| *Control vs Cyclo+TMZ* | 0.81±12.52** | 47.30±4.47* | 72.97±6.75** | 18.20±3.93 | 0.65±23.47*** | 59.68±5.18* | 39.79±2.63 | 10.99±2.93 | 1.07±27.20*** | 40.71±2.61 | 109±6.97** | 38.78±6.22*** |
| *Control vs Cyclo/TMZ* | 0.16±2.56 | 41.00±3.87 | 30.55±3.05 | 5.88±1.16 | 0.02±1.00 | 34.99±2.48 | 38.49±2.54 | 1.70±0.45 | 0.19±4.86* | 29.32±2.10 | 24.77±1.58 | 24.00±4.34* |
| *Control vs TMZ/Cyclo* | 0.77±13.00*** | 32.16±3.28 | 81.06±6.61** | 11.91±1.99 | 0.64±25.63*** | 51.24±4.45* | 0.46±0.03 | 4.90±1.25 | 0.96±24.27*** | 35.43±2.27 | 33.42±1.88 | 34.25±5.49** |
| *Cyclo+TMZ vs Cyclo/TMZ* | 0.65±9.95** | 88.29±7.80** | 103.5±9.58*** | 12.32±2.43 | 0.62±22.47*** | 24.69±1.75 | 78.28±4.72 | 9.27±2.59 | 0.88±22.34*** | 11.39±0.73 | 84.52±5.04* | 14.78±2.67 |
| *Cyclo+TMZ vs TMZ/Cyclo* | 0.04±0.72 | 15.14±1.43 | 8.09±0.62 | 6.29±1.05 | 0.00±0.08 | 8.44±0.73 | 40.25±2.43 | 6.08±1.62 | 0.11±2.93 | 5.27±0.30 | 75.87±4.04 | 4.53±0.72 |
| *Cyclo/TMZ vs TMZ/Cyclo* | 0.60±10.19** | 73.15±6.91** | 111.6±9.11*** | 6.02±0.95 | 0.62±24.54*** | 16.25±1.15 | 38.02±2.29 | 3.19±0.85 | 0.76±19.41*** | 6.10±0.39 | 8.64±0.46 | 10.25±1.85 |

*P< 0.05; **P< 0.01 and ***P< 0.001

**Supplementary Table 6-** Effect of TMZ on GBM cells

|  | GBM 95 | | GBM 02 | | GBM 03 | |
| --- | --- | --- | --- | --- | --- | --- |
|  | *72h* | *144h* | *72h* | *144h* | *72h* | *144h* |
| *Control vs TMZ 100µM* | 14.62±1.36 | 52.92±30.88*** | 6.56±1.70 | 23.32±8.83*** | 17.16±4.23 | 34.61±5.86* |
| *Control vs TMZ 200µM* | 25.44±2.36 | 61.38±35.81*** | 5.27±1.37 | 23.34±8.84*** | 15.72±3.88 | 42.78±7.24** |
| *Control vs TMZ 400µM* | 25.95±2.41 | 69.47±40.53*** | 13.78±3.57 | 41.01±15.54*** | 21.78±5.37* | 57.99±9.81*** |
| *Control vs TMZ 600µM* | 34.52±3.21 | 73.83±43.08*** | 18.03±4.68* | 57.48±21.78*** | 28.80±7.10** | 58.30±9.87*** |
| *TMZ 100µM vs TMZ 200µM* | 10.82±1.00 | 8.45±4.93* | 1.28±0.33 | 0.01±0.00 | 1.43±0.35 | 8.17±1.38 |
| *TMZ 100µM vs TMZ 400µM* | 11.33±1.05 | 16.55±9.65*** | 7.21±1.87 | 17.69±6.70** | 4.61±1.13 | 23.38±3.96 |
| *TMZ 100µMvs TMZ 600µM* | 19.90±1.85 | 20.91±12.20*** | 11.46±2.97 | 34.13±12.94*** | 11.64±2.87 | 23.70±4.01 |
| *TMZ 200µMvs TMZ 400µM* | 0.50±0.04 | 8.08±4.71* | 8.49±2.20 | 17.67±6.69** | 6.05±1.49 | 15.21±2.57 |
| *TMZ 200µM vs TMZ 600µM* | 9.07±0.84 | 12.45±7.26** | 12.75±3.31 | 34.13±12.94*** | 13.08±3.22 | 15.52±2.62 |
| *TMZ 400µM vs TMZ 600µM* | 8.56±0.79 | 4.36±2.54 | 4.25±1.10 | 16.47±6.24** | 7.02±1.73 | 0.31±0.05 |

*P< 0.05; **P< 0.01 and ***P< 0.001
